# Validation of polymorphic Gompertzian model of cancer through *in vitro* and *in vivo* data

**DOI:** 10.1101/2023.04.19.537467

**Authors:** Arina Soboleva, Artem Kaznatcheev, Rachel Cavill, Katharina Schneider, Kateřina Staňková

## Abstract

Mathematical modeling plays an important role in our understanding and targeting therapy resistance mechanisms in cancer. The polymorphic Gompertzian model, analyzed theoretically by Viossat and Noble, describes a heterogeneous cancer population consisting of therapy sensitive and resistant cells. This theoretically promising model has not previously been validated with real-world data. In this study, we provide this validation. We demonstrate that the polymorphic Gompertzian model successfully captures trends in both *in vitro* and *in vivo* data on non-small cell lung cancer (NSCLC) dynamics under treatment. Additionally, for the *in vivo* data of tumor dynamics in patients undergoing treatment, we compare the polymorphic Gompertzian model to the classical oncologic models, which were previously identified as the models that fit this data best. We show that the polymorphic Gompertzian model can successfully capture the U-shape trend in tumor size during cancer relapse, which can not be fitted with the classical oncologic models. In general, the polymorphic Gompertzian model corresponds well to both *in vitro* and *in vivo* real-world data, suggesting it as a candidate for improving the efficacy of cancer therapy, for example through evolutionary/adaptive therapies.

## 1. Introduction

For patients with advanced cancer, an aggressive treatment, aiming at complete cancer eradication, is often ineffective [1, 2]. Instead of pursuing tumor elimination through maximum tolerable dose (MTD), novel therapies – termed evolutionary therapies – aim at anticipating and steering cancer ecoevolutionary dynamics in response to the treatment [2–15]. Mathematical models of cancer’s response to therapy may help us to guide such therapies and, perhaps even more importantly, are needed to understand conditions under which the evolutionary therapies outperform standard of care [5, 16– 25].

Recently, Viossat and Noble [26] analyzed a group of density-dependent polymorphic models assuming two types of cancer cells, therapy-resistant and therapy-sensitive ones, and demonstrated that the containment protocol, which aims to keep the tumor burden below a particular threshold for as long as possible, outperforms standard treatment in all these models in terms of time to progression. Their numerical simulations were performed on one of these models, a two-population Gompertzian model. The model assumes equal growth rates for the sensitive and resistant populations, density-dependent selection, and no cost of resistance, i.e. no assumption that resistant cells are less fit than the sensitive ones when therapy is not applied [14, 27–29]. The model assumes no direct competition between different cell types through competition coefficients, but a shared carrying capacity instead. The absence of the cost of resistance makes the model applicable to a wider range of cancer types, as it was suggested that in some cancers, resistance does not need to have a cost [5, 27, 30, 31].

Viossat and Noble [26] show that in the polymorphic Gompertzian model, adaptive treatment protocols increase time to progression compared to the continuous application of maximum tolerable dose. However, before bringing these models to clinical practice, their ability to fit the real-world data should be evaluated [32, 33]. In this study, we aim to test the polymorphic Gompertzian model’s agreement with *in vitro* data from Kaznatcheev et al. [30] and *in vivo* data from previous clinical studies [34–38] of non-small cell lung cancer (NSCLC) dynamics under therapy.

## 2. Methods

The polymorphic Gompertzian model of cancer analyzed by Viossat and Noble [26] assumes two types of cancer cell populations sensitive and resistant to treatment with sizes *S*(*t*) and *R*(*t*) at time *t*, respectively. The total size of cancer population at time *t* is *N* (*t*) = *S*(*t*) + *R*(*t*). The dynamics of the two populations are described by the ordinary differential equations:

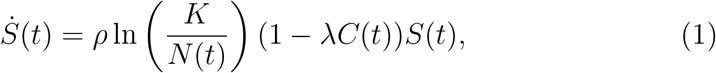

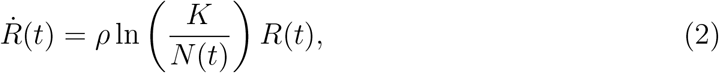

where *C*(*t*) is treatment dosage at time *t, K* is carrying capacity of the tumor (that defines maximum possible size to which the cancer population can grow), *ρ* is growth rate of cancer cells, and *λ* is treatment sensitivity.

Note that competition between sensitive and resistant cells is not explicitly present in the model, besides competing for the space and resources through the carrying capacity. The model also assumes the same growth rate *ρ* for both populations and no cost of resistance.

### 2.1 Fitting the model to in vitro data

We first validated the polymorphic Gompertzian model on *in vitro* data from the study by Kaznatcheev et al. [30]. In their study, two populations of NSCLC cells - sensitive and resistant to treatment - were seeded at different initial proportions with and without addition of cancer-associated fibroblasts (CAF) and the immunotherapy drug Alectinib. In total, the data contains 192 wells: eight different seeding proportions of sensitive cells (0, 0.1, 0.2, 0.4, 0.6, 0.8, 0.9, 1) in four conditions (Drug-, CAF-; Drug-, CAF+; Drug+, CAF-; Drug+, CAF+) and six replicates for each combination. For all wells, sizes of sensitive and resistant populations over time are presented.

We fitted the population dynamics of sensitive and resistant cells for each of the 192 wells to the polymorphic Gompertzian model using the Python GEKKO package [39] with mean squared error as the objective function:

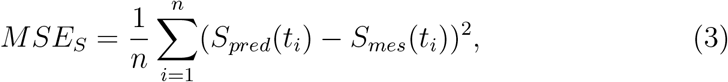

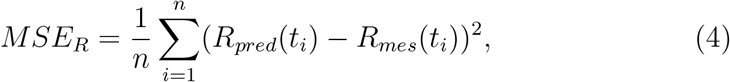

where *S*_*mes*_(*t*_*i*_) and *R*_*mes*_(*t*_*i*_) are measured sizes of sensitive and resistant populations at *i*-th time point, *S*_*pred*_(*t*_*i*_) and *R*_*pred*_(*t*_*i*_) are model-predicted sensitive and resistant population sizes at *i*-th time point, and *n* is the number of time points.

In the fitting procedure, we exploited the explicit treatment modelling through variable *C*(*t*_*i*_), the treatment dosage at time *t*_*i*_. In wells where the drug was not applied, we set *C*(*t*_*i*_) = 0 for all *t*_*i*_. The dynamics in wells with drug were modeled as the treatment variable *C*(*t*_*i*_) set to 0 in time points *t*_*i*_ *<* 20 hours and to 1 at time points *t*_*i*_ ≥ 20 hours. This way, we reflected the conditions of the experiment, where Alectinib was added to the wells 20 hours after seeding [30]. As CAF have a variety of functions and action mechanisms [40], which are challenging for direct modeling, we evaluated its effect implicitly through a comparison of the fitted model’s parameters values (*K, ρ, λ*) in wells with and without CAF. In the wells with seeding proportions 0 and 1 (fully resistant and sensitive wells), we set the measurements of the minor population to zero before fitting the model, as their nonzero measurements present only fluorescent noise. After the polymorphic Gompertzian model was fitted to the wells, the obtained parameter values and their dependence on the initial proportions of sensitive cells and experimental conditions were analyzed.

### 2.2 Fitting the model to in vivo data

We used data from clinical trials of NSCLC patients treated with either immunotherapy drug Atezolizumab or chemotherapy drug Docetaxel [34– 38]. The data contains measurements of the longest diameters of lesions over time, assessed through CT scans. We estimated the volume of the target tumor, following the common assumption that the tumor is a sphere with a diameter equal to the longest measured lesion diameter [41].

Focusing on the tumors with six or more measurements over time, we split the resultant 587 patient cases into five categories based on the trend in the measured data: “Growth”, “Decline”, “Delayed response”, “U-shape” or “Fluctuate” (For details, see Appendix B). We fitted the polymorphic Gompertzian model to the tumor volumes of each of the 587 patients using the Python package GEKKO [39]. We fit the model parameters so that the *MSE* between measured data and the sum of model-predicted sensitive and resistant population sizes defined as

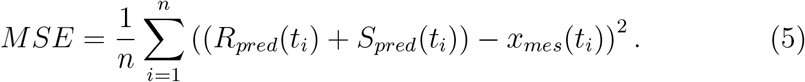

is minimized. In (5), *x*_*mes*_(*t*_*i*_) is measured tumor volume at *i*-th time point, *R*_*pred*_(*t*_*i*_) and *S*_*pred*_(*t*_*i*_) are model-predicted resistant and sensitive population volumes at *i*-th time point, and *n* is number of time points.

As the *in vivo* data contains only total population sizes, initial proportions of sensitive cells were estimated through a grid search (See Appendix A). We modeled the treatment as a constant function *C*(*t*_*i*_) = 1, as in the clinical studies patients received chemotherapy or immunotherapy regularly and in the same dosage [35–38]. After obtaining the best fitting parameters, we analyzed the accuracy of the fit in different trend categories (See Appendix D). We also fitted two best performing classical models of Laleh’s et al.’s study [34] - General Gompertz and General von Bertalanffy models - to the *in vivo* data and compared accuracy of the three models within the trend categories (See Appendix C for more details on the fitting of the classical models and the assessment of the models’ accuracy).

## 3. Results

### 3.1. Validation through in vitro data

#### 3.1.1. The fit of the polymorphic Gompertzian model to data dynamics

We fitted the polymorphic Gompertzian model to *in vitro* dynamics of sensitive and resistant cancer populations. The model’s fit to the measured data in four experimental conditions (Drug-, CAF-; Drug-, CAF+; Drug+, CAF-; Drug+, CAF+) and with seeding proportions of sensitive cells is presented in Fig. 1. Columns represent the experimental conditions of wells. Each row contains wells from one of the intended proportion groups (0.2, 0.4, 0.6, 0.8) corresponding to the seeding proportions of sensitive cells. The actual proportions in wells differ from the intended ones due to experimental variations.

**Figure 1:**
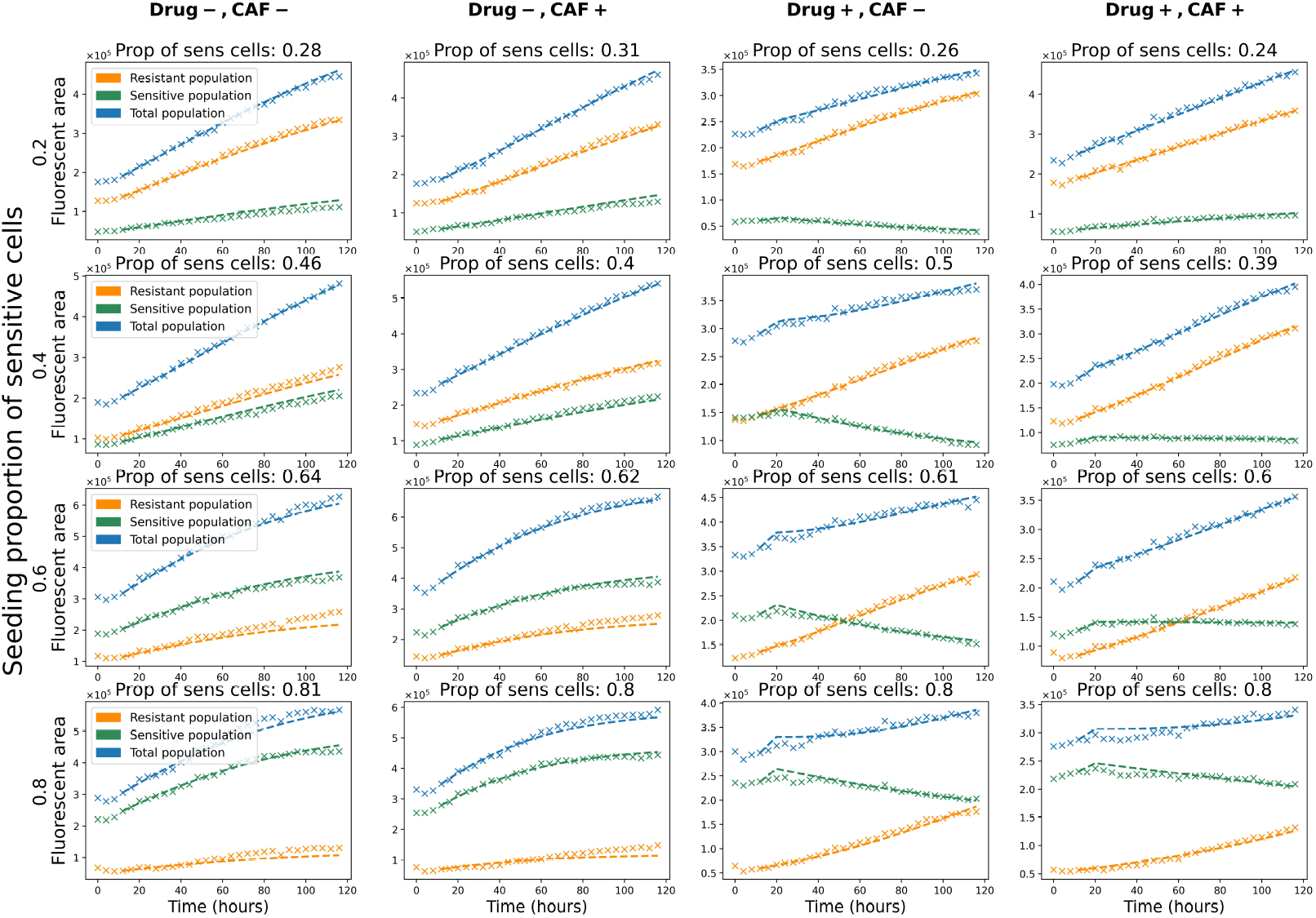
Fit of the polymorphic Gompertzian model to *in vitro* dynamics of sensitive, resistant and total cancer cell populations across four experimental conditions and different initial ratios of sensitive cells. Each graph displays the measured cell counts (crosses) and the model fit (lines) to one well. Sensitive, resistant and total populations are colored in green, orange and blue, respectively. For each well measured proportion of sensitive cells at time point *t*_3_ = 12*h* (start of the model’s fit) is stated above the graph. Columns represent experimental conditions (presence or absence of drug and CAF). Rows contain wells in different conditions with similar initial proportions of sensitive cells.

The trend dynamics were captured well for all experimental and initial conditions. In particular, the model reflected the convex form of the population growth with a slowing of the growth rate closer to a larger size (rows 3 and 4 of Fig. 1). We modeled the therapy start at *t*_*i*_ = 20*h* following the experimental setting and, thus, were able to capture the change of the trend in the sensitive population (columns 3 and 4 of Fig. 1). The population initially was growing, but started to decline once the treatment was added. In the wells with the drug applied (columns 3 and 4 of Fig. 1) the model is able to simultaneously capture the decline of the sensitive population and growth of the resistant population.

In the wells where the sizes of the sensitive and the resistant populations differ significantly, the model fitted the dynamics of the larger population better than the smaller population. This effect is attributed to the common growth rate *ρ* and carrying capacity *K* shared by the sensitive and resistant cancer populations (Eqs. (1)-(2)). The optimization led to a higher weight to the dynamics of a larger population, to decrease the total error. While the relative error of the smaller population increased with the decrease of its proportion, the error for the total population stayed below 5% mean absolute percentage error (*MAPE*) for all proportions (Fig. 2). In monotypic wells (seeding proportions of sensitive cells equal 0 or 1) the error of the model’s fit to the non-seeded population is zero.

**Figure 2:**
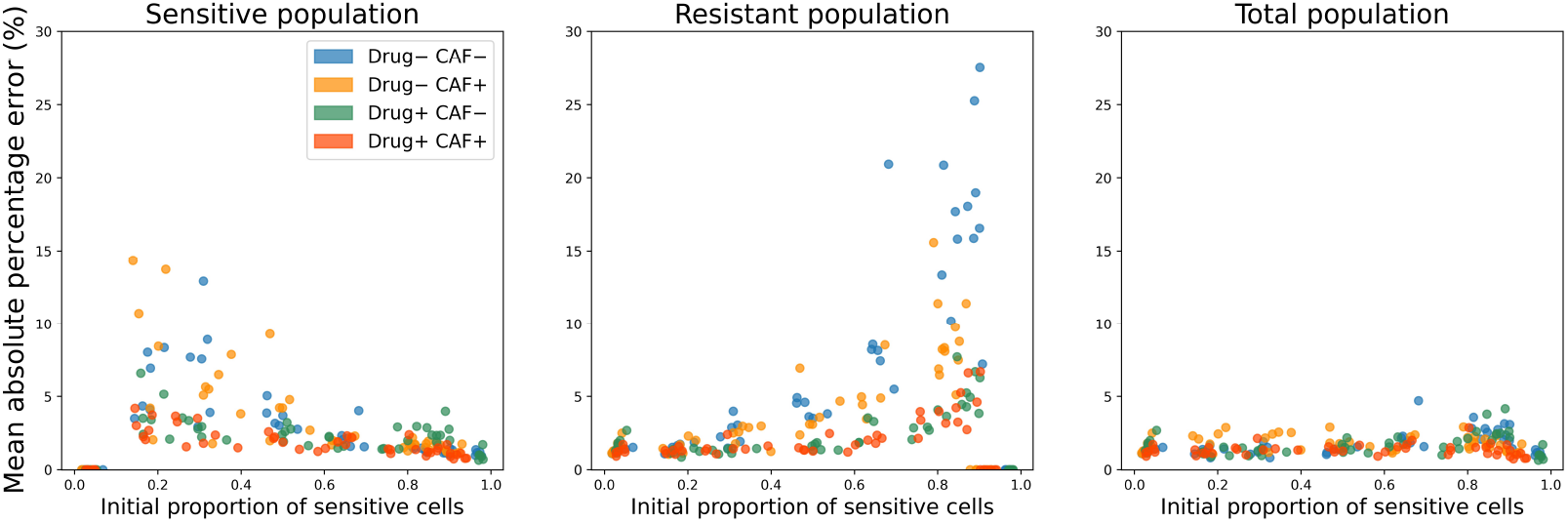
Mean Absolute Percentage Error (MAPE) of the polymorphic Gompertzian model’s fit to *in vitro* dynamics of sensitive, resistant and total cancer cell populations across a range of initial proportion of sensitive cells. Data points are colored according to experimental conditions of the well (presence or absence of drug and CAF).

Fig. 2 also shows that in wells with treatment (Drug+, CAFand Drug+, CAF+), model accuracy was generally higher than in cases without drug. This can be explained by the higher number of parameters in cases with a drug (treatment sensitivity *λ* was fitted only if treatment was present).

#### 3.1.2. Effect of cancer-associated fibroblasts on the model fit

Fig. 3 shows model parameters over wells with different initial ratios of sensitive cells and experimental conditions (presence or absence of drug and CAF). The treatment sensitivity parameter *λ* is significantly smaller in wells where CAF are present, which was confirmed using a Welch’s unequal variance *t*-test (*p*-value= 10^*−*7^). The test repeated with exclusion of the outliers (*S* = 0 or *R* = 0) confirmed the result. Since *λ* represents the magnitude of drug effect on sensitive cells, smaller values of *λ* reflect reduced efficiency of the treatment when CAFs are present.

**Figure 3:**
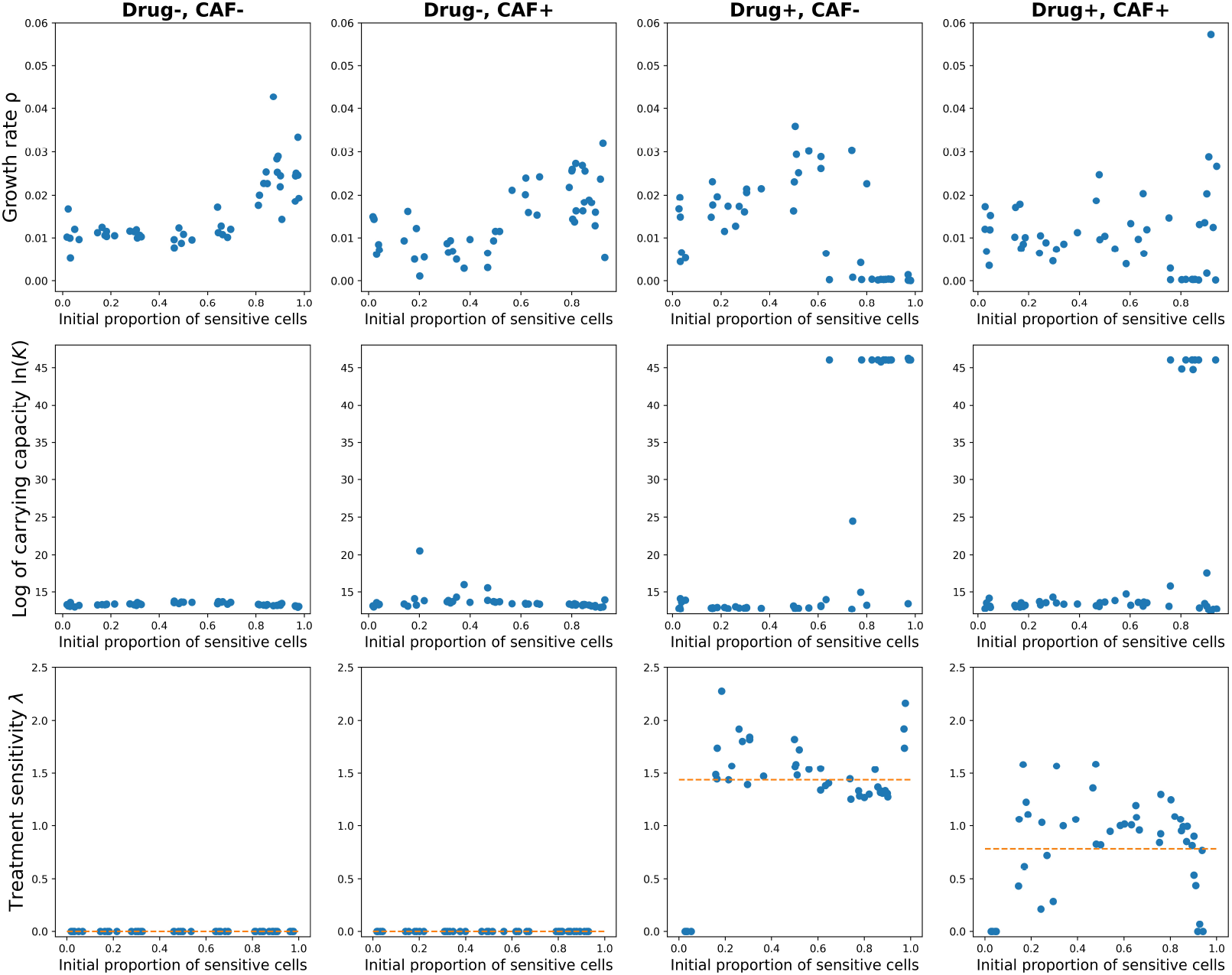
Parameters of the polymorphic Gompertzian model fitted to *in vitro* data across four experimental conditions (indicated in columns) and range of initial proportions of sensitive cells. Each dot in the graphs corresponds to the parameter value of the model fitted to dynamics from one well. The orange line indicates the mean value of treatment sensitivity *λ* in a given experimental condition.

In Fig. 3 we also observe a sudden increase in the fitted carrying capacity *K* and related decay in the fitted growth rate *ρ* for several wells with *p* ≥ 0.6 and drug present. In these cases, a very high *K* value makes the term 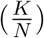 almost constant for the entire measured period (changes in *N* have nearly no effect, as *K* ≫ *N* (*t*), ∀*t*). At the same time, a low growth rate *ρ* compensates for the magnitude of 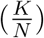. Such parameter values result in an exponential growth for resistant population and an exponential decrease for sensitive population, reflecting the trends observed in the data. We can conclude that for the cases with prevailing sensitive population and drug applied the dynamics switches from Gompertzian to exponential growth/decay.

### 3.2. Validation through in vivo data

#### 3.2.1. The model’s fit to different trend categories

We divided the *in vivo* data into groups based on the displayed trend in tumor growth and evaluated the model’s performance in each of these categories (See Section 2.2). Fig. 4 presents the polymorphic Gompertzian model’s fit to five representative patient cases from each trend category. The two top rows show that the model can accurately capture both “Growth” and “Decline” trends. In these categories, one of the populations has a significantly larger effect on the total population dynamics. In the “Growth” category, the increase of the tumor size is attributed to the proliferation of the resistant population. In the “Decline” category, in contrast, the sensitive population plays a major role, forming the downward trend of the total population size. The polymorphic Gompertzian model is also successful in describing the “U-shape” trend in the data due to incorporated heterogeneity. In this case, sensitive population initially prevails but then decreases under treatment. As the result, resistant population is able to proliferate without competition for space and resources, which leads to regrowth of the tumor.

**Figure 4:**
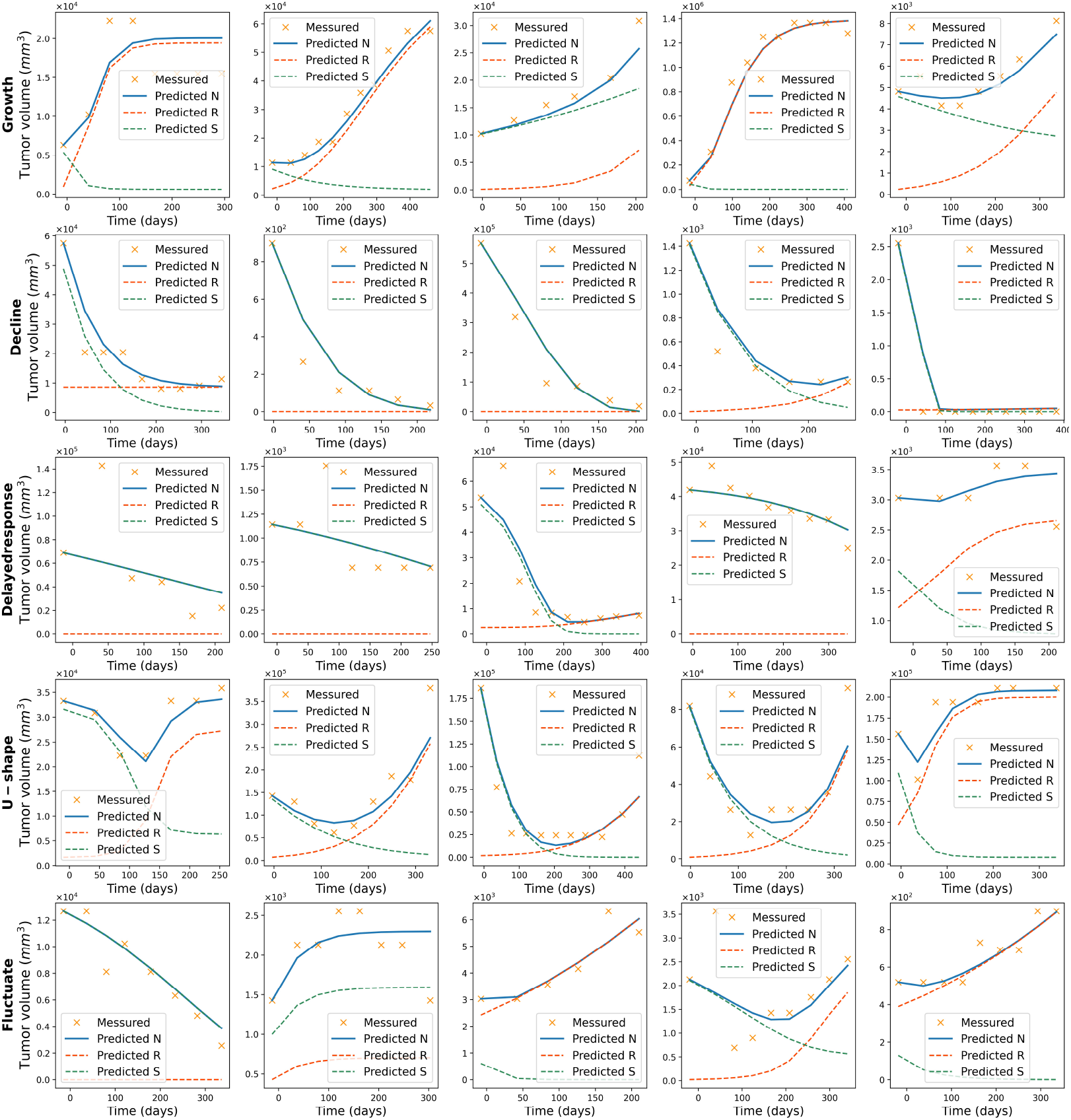
The polymorphic Gompertzian model’s fit to *in vivo* data across five trend categories. Each row shows five patient cases corresponding to one of the trend categories: “Growth”, “Decline”, “Delayed response”, “U-shape” or “Fluctuate”. Measured tumor volume over time is marked with orange crosses. The blue line represents the model’s fit to the total tumor size dynamics, green and orange lines represent predicted dynamics of sensitive and resistant populations sizes respectively.

In the “Delayed response” category, the model is unable to capture the trend. The model tends to ignore initial tumor size increase and fits the data as a monotonic decline. In some cases, where the response is delayed longer, the model describes it as a growth trend and does not capture the subsequent decline (5th graph in the “Delayed response” row of Fig. 4). In the “Fluctuate” category, the polymorphic Gompertzian model estimates data dynamics with a monotonic growth, a monotonic decline, or a U-shape trend. The model is often able to capture the main tendency in the data but ignores smaller fluctuations.

Overall, the lowest mean errors of the model’s fit correspond to “Growth” and “Decline” categories and the highest mean error corresponds to the “Delayed response” trends (See Appendix D). We also conclude that the polymorphic Gompertzian model presents a higher accuracy in the majority of cases comparing to the monomorphic Gompertz and General van Bertalanffy models. The mean error of the polymorphic Gompertzian model’s fit is smaller than that of the General Gompertz model in the “Growth” and “Decline” categories and similar to that of the General von Bertalanffy model. None of the models is capable of describing the “Delayed response” trend and the error values in this category are similar for all three models. The polymorphic Gompertzian model demonstrates a higher accuracy than the other models in the “U-shape” category (See Appendix C for details).

## 4. Discussion

We validated the polymorphic Gompertzian model of Viossat and Noble [26] using *in vitro* and *in vivo* data. Overall, the model fits both studied cases very well. The model assumes (i) a shared carrying capacity between sensitive and resistant cell populations, (ii) the growth rate of the two populations being equal in the absence of drug, and (iii) no competition/cooperation effects in terms of a competition/fitness matrix. Due to these assumptions we were not able to re-evaluate all conclusions of Kazatcheev et al’s game-theoretic model [30].

The polymorphic Gompertzian model may be perceived as implicitly frequency-dependent in *in vitro* case (the growth of the sensitive and resistant populations is dependent on the seeding proportions), as we fitted the model’s parameters separately for different initial proportions of sensitive cells. Nevertheless, the model’s assumptions allowed it to fit both the rich data from *in vitro* experiments and the sparse data from *in vivo* clinical studies.

With the polymorphic Gompertzian model, we took a conceptually different approach to the analysis of the Kaznatcheev et al. [30] *in vitro* data. Kaznatcheev et al. [30] specifically designed a new experimental procedure as a way to measure a competition/fitness matrix [42] that allowed for two distinct fitness functions for the sensitive and resistant cell types. We showed that the polymorphic Gompertzian model could accurately capture the trends in the data with a growth rate *ρ* that is the same for the sensitive and resistant types. We confirmed the anti-treatment effect of cancer-associated fibroblasts. However, we could not confirm or reject Kaznatcheev’s et al. [30] conclusion on strong violations of the cost of resistance (where resistant cells are fitter than sensitive cells even outside treatment).

The future work shall consider extensions of the polymorphic Gompertzian model to cases where sensitive and resistant cells are allowed to have different growth rates and/or when frequency-dependent selection is included in the form of competition matrix. Such extensions can be tested on other existing or proposed *in vitro* datasets that provide data on different cell type proportions [43–46] or population sizes [34, 47, 48].

In this study we demonstrated that the polymorphic Gompertzian model can also be parametrized with the more sparse data from *in vivo* tumor size dynamics of patients undergoing therapy. The polymorphic Gompertzian model provides a good fit of the “Growth” and “Decline” categories, with the error comparable to the error of the best performing classical model of cancer growth – General von Bertalanffy. In the “Growth” category, the polymorphic Gompertzian model described the tumor as consisting mostly of resistant cells. In the “Decline” category, if the tumor size declines monotonically to zero than the model can treat the cancer population as completely sensitive. However, if the last few measurements show a slight increase in the tumor size, the model indicates that the resistant population is still present – allowing for the possibility of relapse and tumor regrowth. The main advantage of the polymorphic Gompertzian model is its ability to capture relapse and regrowth described clinically by the “U-shape” trend – a trend that cannot be captured by any of the six classical models analyzed by Ghaffari Laleh et al. [34].

The polymorphic Gompertzian model (along with the classic models) does not capture the “Delayed Response” trend for *in vivo* data. This could be due to the lack of information on when exactly treatment started: in the clinical studies, the first measurement of the tumor size was usually performed few days before the treatment started. Thus, the initial growth of the tumor in the “Delayed Response” trend might have happened before treatment was first applied. If we had tumor size measurements on the day the therapy started then we could correct for this. We do exactly this correction for Kaznatcheev et al. [30] *in vitro* data to capture a delayed response effect. Alternatively, the *in vivo* “Delayed Response” trend might be attributed to having insufficient concentration of the drug in the tumor at first injection. In this case, data on drug concentration over time would be required for the polymorphic Gompertzian model to describe these cases through a time-dependent treatment variable.

In our study, we demonstrate the ability of the polymorphic Gompertzian model to describe real-world data on cancer under treatment. Going forward, the most promising future development is modelling real-world time dependent treatment, especially adaptive therapy [24, 49, 50]. The “U-shape” trend that the polymorphic Gompertzian model captures in the *in vivo* data presents an undesirable scenario for the patients. In 54 out of 81 cases in the “U-shape” category, the tumor regrows to the initial size or greater. Viossat and Noble’s [26] theoretical analysis of the polymorphic Gompertzian model suggests that for these cases, the implementation of containment therapy could be beneficial and might prolong the time to tumor progression. Contradictory, Kaznatcheev et al. [30] suggest that there would be little benefit to adaptive therapy given the competition/fitness matrices that they estimated *in vitro*. Of course, in the real-world, the behavior of the tumor under containment therapy may differ from the mathematically predictions and, therefore, the effectiveness of such an adaptive therapy should be tested for the cases analyzed in this paper.

## 5. Acknowledgments

This research was supported by the European Union’s Horizon 2020 research and innovation programme under the Marie Sk-lodowska-Curie grant agreement No. 955708 and the Dutch National Foundation projects VI.Vidi.213.139 and OCENW.KLEIN.277.

