## Supplementary for "Validation of polymorphic Gompertzian model of cancer through *in vitro* and *in vivo* data"

### Appendix A. Procedure of fitting the polymorphic Gompertzian model to *in vitro* and *in vivo* data

We fitted the polymorphic Gompertzian model to the *in vitro* and *in vivo* data on cancer population dynamics using GEKKO package. GEKKO is a Python package for machine learning and optimization that allows to dynamically estimate parameters of the differential equations through nonlinear optimization [1].

In the *in vitro* case, we excluded the first three measurements, as they demonstrate fluctuations attributed to the initial stabilization of the system. As the fitting procedure is sensitive to initial values and search bounds, we set physiological lower bounds for the parameters: the growth rate $\rho$ and treatment sensitivity $\lambda$ are assumed positive. We set the lower bound of the carrying capacity $K$ at 0.9 of the maximum measured population size in the well. As the carrying capacity should indicate the growth limit, we assumed it to be bigger than the measured population size at all time, but accounted for the possible 10% measurement error.
To minimize the probability of finding the local minimum instead of the global one, we implemented the grid search over a range of initial parameters values. For more details see GitHub at `https://github.com/SobolevaArina/polymorphic_Gompertzian_model`.

The model was fitted with fixed initial values ($S_{pred}(0) = S_{mes}(0)$, $R_{pred}(0) = R_{mes}(0)$, where $S_{mes}$ and $R_{mes}$ are measured sizes of the sensitive and resistant populations, respectively, and $S_{pred}$ and $R_{pred}$ are sizes of sensitive and resistant populations predicted by the model). We also performed fitting of the model with optimized initial conditions and compared the obtained results. However, we decided to proceed with fixed initial population sizes to keep the number of degrees of freedom low.

The *in vivo* data, unlike the *in vitro* data, does not contain information on the tumor composition. Therefore, initial proportions of sensitive and resistant cells needed to be provided as an input for the optimization procedure. To do that, we implemented an additional grid search for the seeding proportions.

The General Gompertz and General von Bertalanffy models were fitted to the same *in vivo* data, using Python package GEKKO [1]. In the optimization procedure, the MSE between measured tumor volumes and predicted by the model values defined as:

$$MSE = \frac{1}{n}\sum_{i=1}^{n}\left((x_{pred}(t_i) - x_{mes}(t_i)\right)^2. \quad \text{(A.1)}$$

is minimized. In (A.1), $x_{mes}(t_i)$ is measured tumor volume at $i$-th time point, $x_{pred}(t_i)$ is model-predicted volume at $i$-th time point, and $n$ is number of time points. The fittings of the General Gompertz and General von Bertalanffy models were performed with the same procedure as for the polymorphic Gompertzian model. For more details on the fitting procedure see GitHub (`https://github.com/SobolevaArina/polymorphic_Gompertzian_model`).

### Appendix B. Trend categories in tumor volume dynamics in *in vivo* data

*In vivo* data of 587 patients was split into five categories based on the observed tumor volume dynamics. For each patient, we calculated a vector of tumor volume differences between measurmets made at time point $t_{i+1}$ and at time point $t_i$ for all $i$. If the difference is positive, then the tumor volume increases between measurements and vice versa. Based on values in the difference vector, we categorized patient cases into trend groups. Categories, the number of cases in each of them, and criteria for classification of the case into one of the categories are presented in Table B.1.

| Category | Number of cases | Criteria |
|---|---|---|
| Growth | 98 | All differences are positive or the sum of positive differences is at least two times greater than the absolute sum of negative differences |
| Decline | 239 | All differences are negative or the absolute sum of negative differences is at least two times greater than the sum of positive differences |
| Delayed response | 65 | First non-zero difference value is positive, the difference vector contains a negative value and the last measurement is smaller than the first one |
| U-shape | 81 | Negative difference value followed by positive, but not vice versa, fluctuations of 10% excluded |
| Fluctuate | 104 | All others |

Table B.1: Categories of patients according to the observed trend in tumor volume dynamics, the number of cases in each category and criteria of classification.

### Appendix C. Comparison of the polymorphic Gompertzian model's fit with the fits of the General Gompertz and General von Bertalanffy models

The fit of the polymorphic Gompertzian model to *in vivo* data was compared to the fit of monomorphic General Gompertz and General von Bertalanffy models fitted with the same procedure. These models previously demonstrated the highest accuracy compared to the other textbook models, when fitted to this dataset [2]. The models' equations are as follows:

General Gompertz:

$$\dot{V}(t) = V^{\mu}(\delta - \gamma \ln V), \tag{C.1}$$

where $V$ is size of the tumor in $mm^3$, $\gamma$ is the maximum net growth rate of cancer population, $\mu$ and $\delta$ are constants.

General von Bertalanffy:

$$\dot{V}(t) = \alpha V^{\mu} - \beta V, \tag{C.2}$$

where $V$ is size of the tumor in $mm^3$, $\alpha$ is the birth rate of cancer cells, $\beta$ is the death rate of cancer cells and $\mu$ is a constant.

The General Gompertz and General von Bertalanffy models were fitted to the *in vivo* data using Python package GEKKO [1]. Fig. C.1 demonstrates fits of the three models to the five representative example cases from "Growth", "Decline", "Delayed response", "U-shape" and "Fluctuate" categories. All three models can describe "Growth" and "Decline" categories well with the models' dynamics similar to each other (Rows $1-2$ of Fig. C.1). None of the models is able to capture the "Delayed response" trend. In the "U-shape" category the polymorphic Gompertzian model outperforms both General Gompertz and General von Bertalanffy models.

We also compared errors of the polymorphic Gompertzian, General Gompertz and General von Bertalanffy models. We calculated normalized mean square error $nMSE$, which is a mean squared error divided by square of the largest measured volume in the case. For the polymorphic Gompertzian model:

$$nMSE = \frac{1}{n}\sum_{i=1}^{n}\left(\frac{S_{pred}(t_i) + R_{pred}(t_i) - x_{mes}(t_i)}{x_{mes}^{max}}\right)^2, \tag{C.3}$$

with $x_{mes}(t_i)$ being the measured tumor volume at the $i$-th time point, $R_{pred}(t_i)$ and $S_{pred}(t_i)$ the resistant and sensitive population volumes at the $i$-th time point predicted by the model, $x_{mes}^{max}$ the maximum measured tumor volume and $n$ the number of measurements.

For General Gompertz and General von Bertalanffy models:

$$nMSE = \frac{1}{n}\sum_{i=1}^{n}\left(\frac{x_{pred}(t_i) - x_{mes}(t_i)}{x_{mes}^{max}}\right)^2, \tag{C.4}$$

where $x_{mes}(t_i)$ - measured tumor volume at $i$-th time point, $x_{pred}(t_i)$- tumor volume at $i$-th time point predicted by the model, $x_{mes}^{max}$ - maximum measured tumor volume, $n$ - number of measurements.

Violin plots of the $nMSE$ distribution for the General Gompertz, General von Bertalanffy and the polymorphic Gompertzian models across five trend categories are presented in Fig. C.2. The $nMSE$ of the polymorphic Gompertzian model's fit is lower than the one of the General Gompertz model's fit in the "Growth" and "Decline" categories and similar to the fit of the General von Bertalanffy model. The polymorphic Gompertzian model has a higher accuracy than the other models in the "U-shape" category. Table C.1 presents $p$-values of $t$-test between the models' $nMSE$ in trend categories.

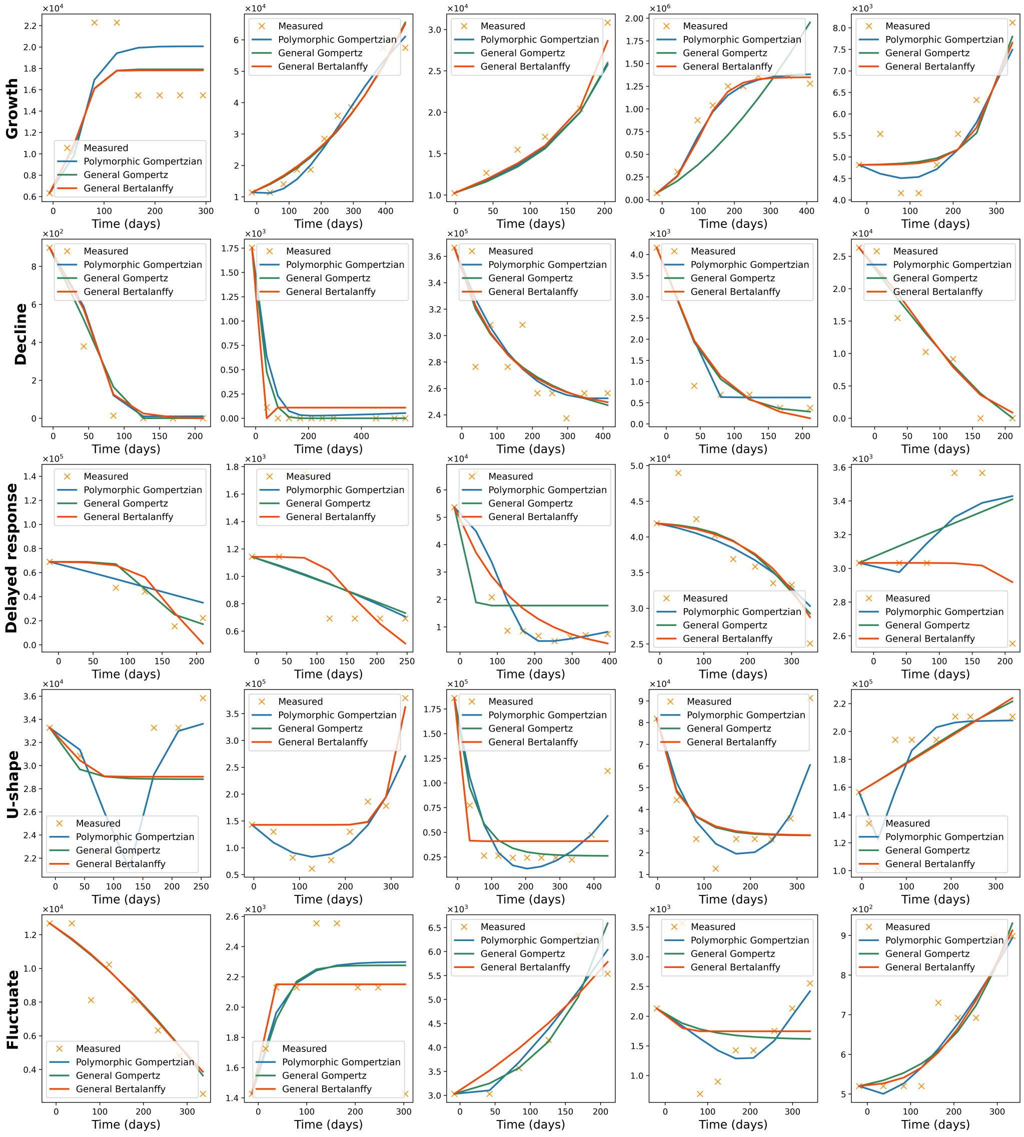


Figure C.1: Fits of the polymorphic Gompertzian, General Gompertz, and General von Bertalanffy models to *in vivo* data for five patient cases in each trend category: “Growth”, “Decline”, “Delayed response”, “U-shape” and “Fluctuate”. Data points are marked with orange crosses. Blue, green, and red lines demonstrate fits of polymorphic Gompertzian, General Gompertz, and General von Bertalanffy models respectively.

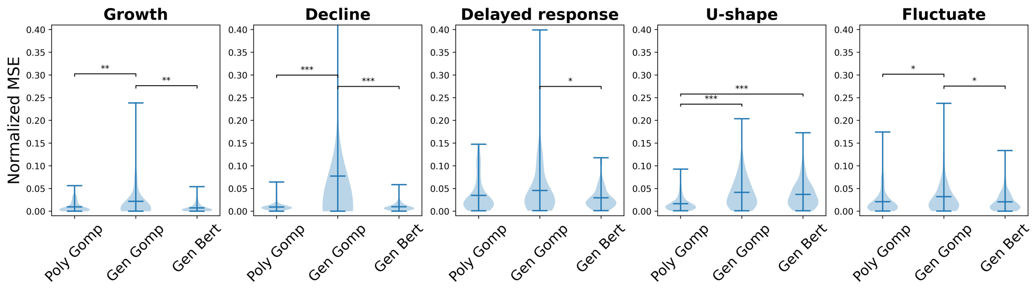


Figure C.2: Distributions of nMSE of the polymorphic Gompertzian, General Gompertz and General von Bertalanffy models' fits to *in vivo* data from five trend categories ("Growth", "Decline", "Delayed response", "U-shape" or "Fluctuate"). In the "U-shape" category nMSE of the polymorphic Gompertzian model is significantly lower than of the other two models. Stars denote statistically significant differences between means of two groups ($\star$ - $p$-value $< 0.05$, $\star\star$ - $p$-value $< 0.01$, $\star\star\star$ - $p$-value $< 0.001$).

| | | **Poly Gomp** | **Gen Gomp** | **Gen Bert** |
|---|---|---|---|---|
| Growth | Poly Gomp | 1 | | |
| | Gen Gomp | **0.007** | 1 | |
| | Gen Bert | 0.1 | **0.001** | 1 |
| Decline | Poly Gomp | 1 | | |
| | Gen Gomp | $\mathbf{2 \cdot 10^{-6}}$ | 1 | |
| | Gen Bert | 0.5 | $\mathbf{2 \cdot 10^{-6}}$ | 1 |
| Del Resp | Poly Gomp | 1 | | |
| | Gen Gomp | 0.2 | 1 | |
| | Gen Bert | 0.4 | **0.04** | 1 |
| U-shape | Poly Gomp | 1 | | |
| | Gen Gomp | $\mathbf{9 \cdot 10^{-8}}$ | 1 | |
| | Gen Bert | $\mathbf{7 \cdot 10^{-7}}$ | 0.4 | 1 |
| Fluctuate | Poly Gomp | 1 | | |
| | Gen Gomp | **0.02** | 1 | |
| | Gen Bert | 1 | **0.01** | 1 |

Table C.1: Comparison of the models' fit errors in trend categories. The table presents $p$-values of $t$-tests between two groups of $nMSE$. Bold text indicates statistically significant $p$-values.

### Appendix D. Parameter values of the polymorphic Gompertzian model across trend categories

We assessed the fitted parameter values of the polymorphic Gompertzian model across trend categories. The distributions of the fitted carrying capacity $K$, growth rate $\rho$, treatment sensitivity $\lambda$, and the estimated initial

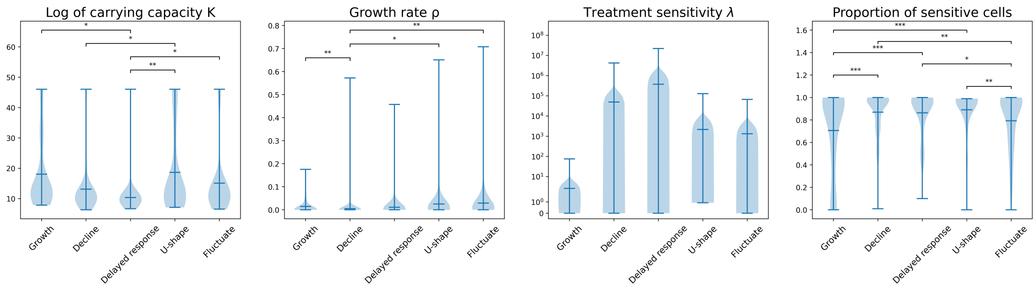


Figure D.1: Violin plots for the logarithmic values of fitted carrying capacity $K$, growth rate $\rho$, treatment sensitivity $\lambda$ and estimated values of initial proportions of sensitive cells of the polymorphic Gompertzian mode's fit to *in vivo* data across five trend categories("Growth", "Decline", "Delayed response", "U-shape", and "Fluctuate"). Stars denote statistically significant differences between means of two groups ($\star$ - $p$-value $< 0.05$, $\star\star$ - $p$-value $< 0.01$, $\star\star\star$ - $p$-value $< 0.001$).

proportions of sensitive cells are presented in Fig. D.1. To ensure the statistical significance of the differences between groups, we implemented $t$-test.

We observed that the distributions of the model parameters and the initial proportion of sensitive cells across the trend categories are in agreement with the biological features of these groups (Fig. D.1). The carrying capacity captures restrictions of physical space, availability of required nutrients, and also implicitly includes other environmental factors limiting cells' growth, such as the immunological response of the organism to the tumor growth [3]. The highest carrying fitted capacity $K$ is observed in "U-shape" category. It emphasizes that the carrying capacity does not affect the form of dynamics, because the tumor does not grow to the size, at which it would slow its growth rate. In some cases from this category the obtained carrying capacity was at the maximum value possible in the coding environment ($10^{45}$). That may indicate the absence of the carrying capacity and, thus, exponential growth of the cancer populations. In the "Growth" category the carrying capacity values are also elevated. This way the model reflects that the tumor has sufficient resources and space for the growth. In "Delayed response" and "Decline" categories $K$ is significantly lower, indicating restrictions of the limited space and resources on the tumor.

The distributions of parameters in the "Delayed response" and the "Decline" categories are similar (Fig. D.1), as the model does not capture the initial increase in the size and instead describes the "Delayed response" trend as a monotonic decline. In the "Decline" category the growth rate $\rho$ is lower and the treatment sensitivity $\lambda$ is higher than in all others categories, except for the "Delayed response". This indicates that the tumors from "Decline" and "Delayed response" categories grow slowly in the absence of drug and have strong response to the applied treatment.

### Appendix E. Exponential growth and decay trends in the polymorphic Gompertzian model

When the polymorphic Gompertzian model was fitted to *in vitro* data, we observed a sudden 'jump' in the obtained carrying capacity values (Fig. 3, main text). In several cases with higher proportions of sensitive cells ($p \geq 0.6$) and drug present, the value of the fitted carrying capacity $K$ was close to $10^{45}$, which is the maximum possible number in the coding environment.

After conducting additional tests, we established that the optimization procedure attempts to set the carrying capacity to the maximum possible value in the given search space. We hypothesize that this way the model indicates the absence of the carrying capacity and, hence, exponential growth of the resistant population and exponential decay of the sensitive population (as in the cases with drug present, the sensitive population declines).

As the model with shared growth rates fits the larger population better, the absence of the carrying capacity is likely attributed to the exponential decrease of the sensitive population. It is also justified by the fact that for some wells which contained only the sensitive population, the obtained carrying capacity was also at the highest possible number in the coding environment. If the resistant population is present in the well with the high $K$, it is small, and, thus, has enough space and resources to grow. It is not clear whether the growth is exponential or logistic at the initial stage of increasing growth.

The growth can be both exponential or logistic (at the initial stage), as both cases can be fitted well with the polymorphic Gompertzian model with the high carrying capacity. The extension of the model with the separate growth rates and/or separate contributions to the carrying capacity for the sensitive and resistant populations can allow fitting trends in both populations better and assess the reasons for the switch from Gompertzian growth to exponential.

Carrying capacity values equal to the maximum number in the coding environment were also observed in several *in vivo* cases. Measurements in such cases coincide with exponential growth, which the model accurately fits.

These cases require special attention, as the presence of the carrying capacity and/or competition between treatment-sensitive and treatment-resistant populations lead to better outcomes of evolutionary therapy, which aims to anticipate and steer the evolution of resistance in the cancer population. Thus, patients with exponential tumor growth would likely benefit only little from the adaptive therapy. However, for the majority of the patients, the carrying capacity is finite, and according to Viossat and Noble's theoretical conclusions, they would benefit more from the containment approach than from the MTD approach in terms of time to progression [4].
